# UCP1 Mitigates Hepatic Steatosis and Fibrosis Independent of Cold Exposure

**DOI:** 10.1101/2025.08.05.666925

**Authors:** John A. Haley, Johnny Le, Rui Li, Ekaterina D. Korobkina, Flaviane De Fatima Silva, Maria Gaughan, Timothy P. Fitzgibbons, Nuria Martinez, Qingbo Chen, Shelagh M. Fluharty, Huawei Li, Caroline A. Lewis, John E. Harris, Lihua Julie Zhu, Alan C. Mullen, Cholsoon Jang, David A. Guertin

## Abstract

Non-shivering thermogenesis by brown adipose tissue (BAT) is a promising target for anti-obesity therapies, making its regulatory mechanisms of significant translational interest. While cold-induced BAT thermogenesis (CIT) is well characterized, certain high-calorie diets can also activate BAT in the absence of cold, a process known as diet-induced thermogenesis (DIT). Despite its potential relevance to modern human diets and lifestyles, the mechanisms and physiological relevance underlying DIT remain poorly understood. Here, we show that DIT reduces adiposity and protects against hepatic steatosis and fibrosis in males but not female mice. Moreover, adipose tissue-specific ablation of uncoupling protein 1 (UCP1) reveals that BAT is the primary mediator of DIT but that non-adipocyte UCP1 also contributes to body weight regulation. Transcriptome analysis suggests that DIT is triggered by intrinsic metabolic stress, distinguishing it from CIT, which is driven by sympathetic tone. Finally, BAT-specific arteriovenous metabolomics identifies glucose as the predominant circulating fuel for DIT. These findings uncover distinct molecular and metabolic features of DIT, highlighting opportunities to harness BAT activity for treating obesity and metabolic diseases without requiring cold exposure.

## Introduction

Obesity impacts over 1 billion people worldwide and is a major risk factor for chronic diseases, including type 2 diabetes, cardiometabolic diseases, neurological disorders, and cancer. Metabolic dysfunction-associated steatotic liver disease (MASLD)^1^ is linked to obesity^2^ and progresses to metabolic dysfunction-associated steatohepatitis (MASH), a severe condition associated with irreversible fibrosis, cirrhosis, and hepatocellular carcinoma (HCC). Accurately modeling MASLD/MASH^3^ remains a challenge and therapeutic options are limited.

In adult humans, the presence of brown adipose tissue (BAT) is associated with resistance to cardiometabolic diseases, including MASLD^4^, but functional connections remain elusive. BAT is a specialized type of adipose tissue that expends calories to generate body heat during cold exposure. This process, also known as non-shivering thermogenesis (NST), is distinguished from shivering thermogenesis by skeletal muscle and is stimulated by the sympathetic nervous system. NST is driven by the inner mitochondrial membrane protein uncoupling protein 1 (UCP1), which facilitates proton leak back into the mitochondrial matrix, thereby uncoupling ATP synthesis from the electron transport system and causing energy to be lost as heat^5^. Stimulating NST to enhance energy expenditure is under investigation as a therapeutic strategy to combat obesity and metabolic diseases.

In contrast to cold exposure, certain high-fat, cafeteria-style diets can also trigger BAT thermogenesis in the absence of cold stress, called diet-induced thermogenesis (DIT). This poorly understood phenomenon, first described in 1979^6^, occurs at thermoneutrality (a range around 30°C for mice), and is associated with UCP1 induction^7–9^. DIT is also distinct from the immediate postprandial increase in body heat production. The most common diet used to stimulate DIT in mice is a 45% fat/17% surcorse diet^7–9^ and several studies using whole-body *Ucp1* knockout (*Ucp1-/-)* mice reported that DIT increases energy expenditure and protects against adiposity^7,10–13^. These studies also showed that whole-body *Ucp1-*deficient mice living at thermoneutrality gain more weight even when eating a control (low fat) diet^7^, collectively arguing that UCP1 activity contributes to the adiposity set point at thermoneutrality regardless of the diet. These findings led to the hypothesis that a deficiency in UCP1 could promote obesity, and that UCP1 thus plays a role in defending the body against excess nutrients, in addition to its classic role in maintaining body temperature in a cold environment^14–16^. However, this idea has been met with skepticism, as it is argued that evolutionary pressure for a mechanism protecting against diet-induced obesity is unlikely to have existed^9,17^. Moreover, not all commercially available diets reliably elicit DIT, suggesting that a specific nutrient combination, rather than the caloric content of the diet alone, is necessary^9,16^. Interestingly, recent studies suggest that UCP1 may be expressed in some non-adipocyte cell populations, raising additional questions about the BAT-specific role of UCP1 in this process, as all previous studies of UCP1’s role in DIT used the whole-body UCP1 knockout model^18–20^. Nevertheless, the potential for BAT to influence energy balance and obesity in humans living at thermoneutrality remains an important and open area of investigation.

In this study, we employed both a whole-body *Ucp1* knockout (*Ucp1 -/-*) and a conditional adipose-tissue-specific *Ucp1* knockout model (*adiponectin-Cre;Ucp1*) to investigate whether UCP1 regulates liver health in the absence of thermal stress. We found that prolonged consumption of the 45% fat/17% sucrose diet induces UCP1 in the BAT of both in males and females, but that UCP1 only protects males from diet-induced adiposity, hepatic inflammation, and fibrosis, uncovering a sex-dependent function for UCP1 in modulating liver disease in the absence of cold stress. Interestingly, we also provide evidence that non-adipose tissue UCP1 contributes to body weight regulation in thermoneutral-housed mice, supporting a previously unrecognized role for non-adipocyte UCP1 in energy balance. We also found that DIT exhibits a molecular signature consistent with a metabolic stress response, distinguishing it from cold-induced thermogenesis (CIT), which is well-known to be driven by sympathetic tone. Yet despite the wide divergence in molecular signatures between cold- and diet-induced thermogenesis, DIT, like CIT, utilizes circulating glucose as a primary fuel source, suggesting a link between UCP1 activity and glucose dependency. These findings advance a fundamental understanding of UCP1 functions that are independent of cold stress, potentially opening new avenues for therapeutic intervention aimed at treating obesity and chronic metabolic diseases.

## Results

### Diet-induced UCP1 Expression in BAT Precedes Hepatomegaly and Glucose Intolerance

To investigate BAT metabolism during DIT and its impact on health, we established a DIT protocol by adapting 6-week-old male mice to thermoneutrality (30°C) for 4 weeks, which inactivates non-shivering thermogenesis. Given that not all commercially available diets reliably elicit DIT^9^, we optimized a dietary intervention protocol; mice are first adapted to thermoneutrality for 4 weeks then fed a high-fat diet (HFD) similar in caloric distribution to a fat-rich human diet that was 45% fat, 17% sucrose, and 20% protein^21^ or a calorically matched low-fat diet (LFD) that was 10% fat, 17% sucrose, and 20% protein, where the high fat content was replaced by a mix of complex carbohydrates that resemble the mix of a standard chow diet.

In male mice after 8 or 16 weeks of the diet provision, HFD feeding resulted in higher levels of UCP1 protein expression and an increased abundance of multi-locular brown adipocytes characteristic of active BAT [Extended Fig. 1a, 1b]. In contrast, mice on LFD had brown adipocytes that appear more like white adipocytes (i.e. contain a single large lipid droplet) characteristic of mice in the absence of cold stress. After 8 weeks of feeding, mice on the HFD were heavier than mice on LFD due to increased white adipose tissue (WAT; subcutaneous (SAT) and visceral adipose tissue (VAT)) expansion [Extended Fig. 1c, 1d], which was at least partly due to adipocyte hypertrophy [Extended Fig. 1e]. Total liver mass and glucose tolerance were unchanged at the 8-week time point [Extended Fig. 1d, 1f]. After 16 weeks of feeding, mice on HFD not only had increased body mass and adiposity, but their livers nearly doubled in size [Extended Fig. 1g, 1h] and had severe steatosis [Extended Fig. 1i]. These mice also became more glucose intolerant [Extended Fig. 1j]. Thus, HFD feeding under thermoneutrality induces UCP1 and the formation of multi-locular adipocytes in BAT and the expansion of WAT by 8 weeks. Importantly, for studies appearing later in this report, these changes precede the development of hepatomegaly and glucose intolerance. Having established a robust DIT assay, we next investigated whether DIT might also protect against liver disease.

### UCP1 Suppresses Diet-induced MASH and Fibrosis Independent of Cold Exposure

While UCP1’s role in regulating adiposity in thermoneutral conditions is established, its impact on the liver is unknown. To investigate this, we first subjected male whole-body UCP1 knockout (*Ucp1^-/-^*) mice and their littermate wild-type (WT) controls to the LFD versus HFD feeding protocol at thermoneutrality for 16 weeks [Fig. 1a]. Total body mass [Fig. 1b], BAT mass [Fig. 1c], and liver mass [Fig. 1d] were all substantially enlarged in the HFD *Ucp1^-/-^* group versus HFD WT, with liver mass increasing by 62.5% in the HFD *Ucp1^-/-^* group relative to HFD WT. Subcutaneous white adipose tissue (SAT) mass was also significantly larger in the HFD *Ucp1^-/-^* group [Fig. 1e], while visceral adipose tissue (VAT) mass was significantly decreased compared to HFD WT [Fig. 1f], indicating depot selective effects. Food intake was unchanged [Fig. 1g] while metabolic efficiency (i.e. the ability to convert energy from food into stored fat) increased in *Ucp1^-/-^* mice, consistent with previous studies^7,10^ [Fig. 1h]. Individual brown adipocytes in the HFD *Ucp1^-/-^* group also lost their multi-locular appearance [Fig. 1i] while the white adipocytes appeared to be similar in size between *Ucp1^-/-^* and WT mice regardless of the diet [Fig. 1i]. These adipose tissue phenotypes align with previous studies, while hepatomegaly is a previously unrecognized consequence of whole-body *Ucp1* ablation during high-fat feeding at thermoneutrality.

**Figure 1.**
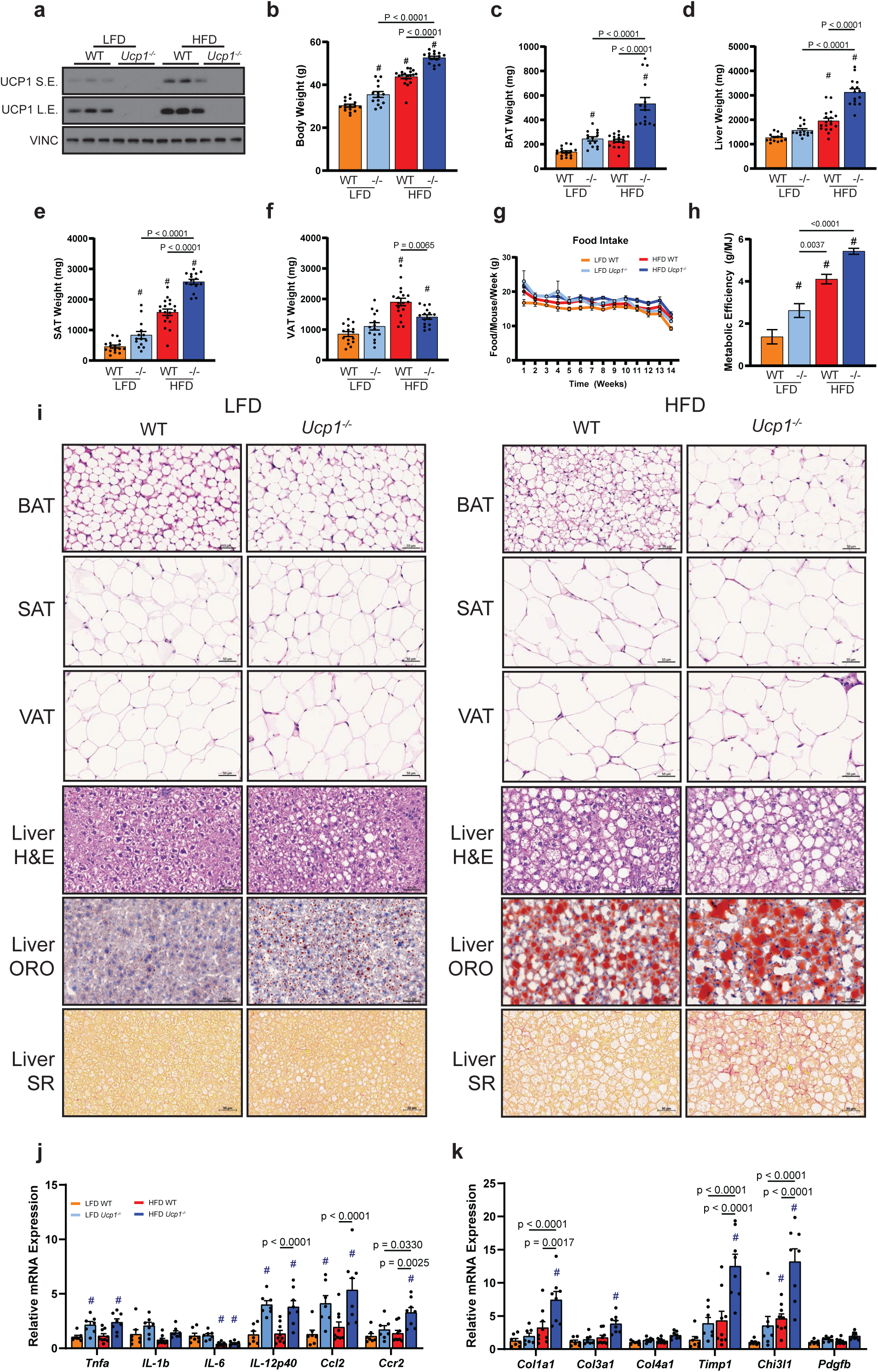
HFD-fed male mice develop liver fibrosis in the absence of UCP1. **a**, Immunoblot of BAT from male WT and UCP1^-/-^ mice fed LFD or HFD for 16-weeks. **b-f**, Body weight (**b**), BAT weight (**c**), Liver weight (**d**), SAT and VAT weights (**e,f**) from male WT and UCP1^-/-^ mice fed LFD or HFD for 16 weeks. Data are mean ± s.e. P values were determined via one-way ANOVA with Tukey’s multiple comparisons, where #, denotes P<0.05 compared to WT LFD. **g**, Food intake from male WT and UCP1^-/-^ mice fed LFD or HFD. **h**, Metabolic efficiency calculated by the ratio of body weight gain vs MJ of food consumed from male WT and UCP1^-/-^ mice fed LFD or HFD. Data are mean ± s.e. P values were determined via one-way ANOVA with Tukey’s multiple comparisons, where #, denotes P<0.05 compared to WT LFD. **i**, Hematoxylin and Eosin (H&E) staining of BAT, SAT, VAT, and liver, as well as oil red-O and Sirius red staining of the liver from male WT and UCP1^-/-^ mice fed LFD or HFD for 16 weeks. **j,k**, RT-qPCR of livers looking at inflammation makers (**j**) and fibrosis markers (**k**) from male WT and UCP1^-/-^ mice fed LFD or HFD for 16 weeks. Data are mean ± s.e. P values were determined via one-way ANOVA with Tukey’s multiple comparisons, where #, denotes P<0.05 compared to WT LFD.

The increased liver mass in HFD *Ucp1^-/-^* group is at least partly attributable to increased lipid droplet size, indicated by H&E and Oil Red O stain [Fig. 1i]. Concomitantly, several inflammation markers were elevated in the liver of HFD *Ucp1^-/-^* mice [Fig. 1j]. Notably, hepatic inflammation was also moderately elevated in LFD *Ucp1^-/-^* mice [Fig. 1j], which was associated with increased macrophages, including Kupffer cells, monocyte-derived macrophages, and capsular macrophages [Extended Fig. 2a-2e], but not other immune cells [Extended Fig. 2f-2i]. Portal circulation cytokines were also elevated in HFD *Ucp1^-/-^* [Extended Fig. 2j-2l]. Sirius Red staining further revealed severe hepatic fibrosis in HFD *Ucp1^-/-^* mice [Fig. 1i] compared to the other groups. Consistently, the livers of HFD *Ucp1^-/-^* had elevated expression of the fibrosis marker genes, including *Col1a1, Col3a1, Timp1*, and *Chi3l1* [Fig. 1k]. These observations were striking because the same diet takes over 40 weeks to cause only mild hepatic fibrosis in WT mice housed at room temperature^3^. Overall, these data reveal a hitherto unappreciated cold-independent role for whole-body UCP1 in resisting diet-induced hepatic steatosis, inflammation and fibrosis.

### UCP1 Loss Does Not Exacerbate Hepatic Inflammation and Fibrosis in Females during DIT

We next examined whether DIT-associated UCP1 induction also plays a protective role in females. After 16 weeks of HFD feeding, female BAT also exhibits increased UCP1 expression [Fig. 2a]. However, total body weight did not differ between *Ucp1^-/-^* and WT females on either LFD or HFD [Fig. 2b]. While HFD *Ucp1^-/-^* females did show increased total BAT mass relative to HFD WT [Fig. 2c], there were no differences in the mass of liver [Fig. 2d], SAT [Fig. 2e], or VAT [Fig. 2f] between the two groups on either diet. HFD *Ucp1^-/-^* females showed no significant differences in food intake or metabolic efficiency from WT in either diet group [Fig. 2g, 2h]. Interestingly, unlike *Ucp1^-/-^* males, which developed profound BAT whitening on HFD [Fig. 1i], *Ucp1^-/-^* females on HFD maintained a multi-locular morphology characteristic of active brown adipocytes [Fig. 2i]. Moreover, while livers in *Ucp1-/-* females showed increased hepatic steatosis compared to WT on both the LFD and HFD [Fig. 2i], the increase in steatosis was not associated with elevated inflammation [Fig. 2j] or fibrosis [Fig. 2k]. Thus, at least under these conditions, UCP1 loss does not exacerbate liver disease in female mice, suggesting the presence of sexually dimorphic mechanisms.

**Figure 2.**
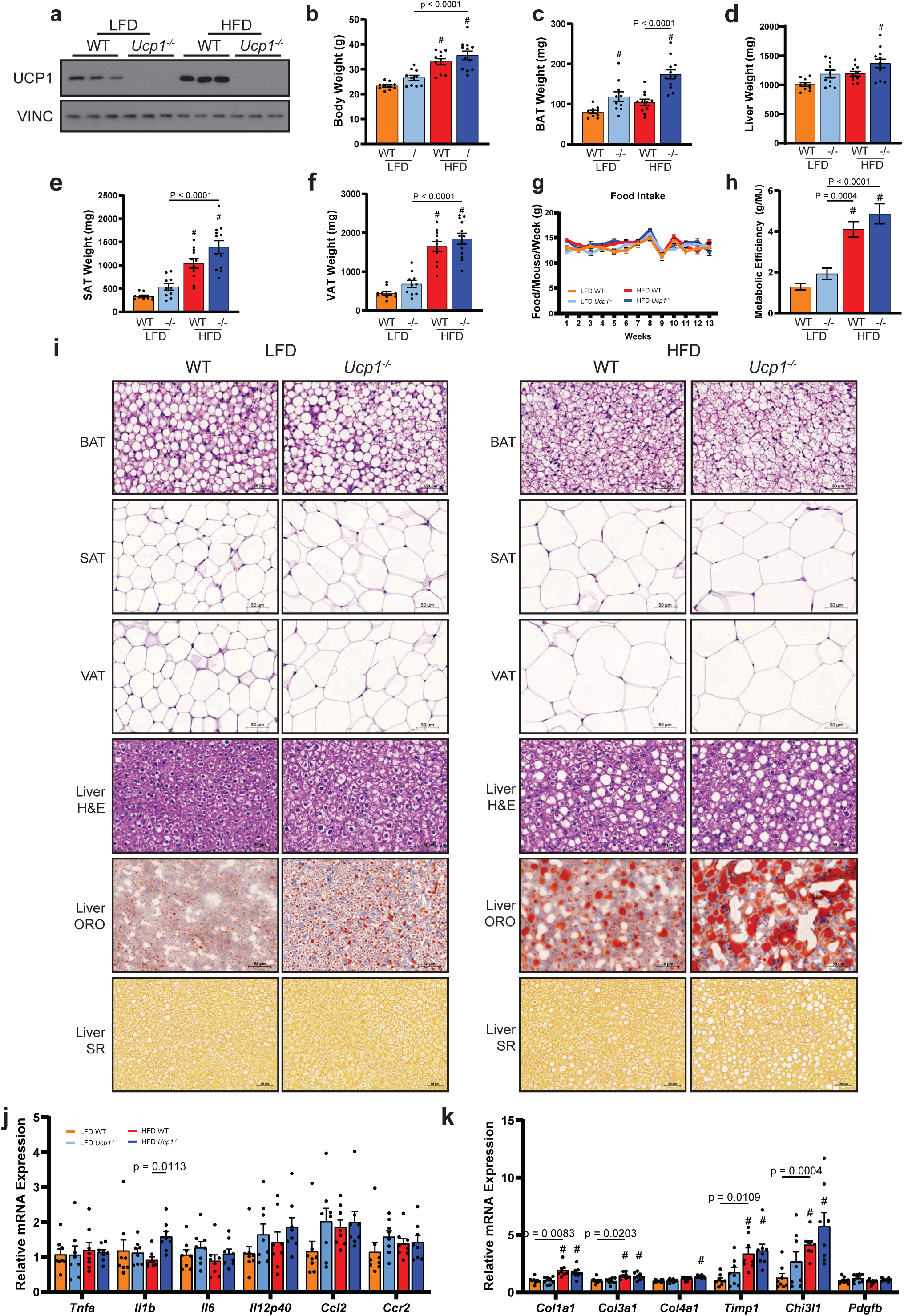
HFD-fed female mice do not develop liver fibrosis in the absence of UCP1. **a**, Immunoblot of BAT from female WT and UCP1^-/-^ mice fed LFD or HFD for 16-weeks. **b-f**, Body weight (**b**), BAT weight (**c**), Liver weight (**d**), SAT and VAT weights (**e,f**) from female WT and UCP1^-/-^ mice fed LFD or HFD for 16 weeks. Data are mean ± s.e. P values were determined via one-way ANOVA with Tukey’s multiple comparisons, where #, denotes P<0.05 compared to WT LFD. **g**, Food intake from female WT and UCP1^-/-^ mice fed LFD or HFD. **h**, Metabolic efficiency calculated by the ratio of body weight gain vs MJ of food consumed from female WT and UCP1^-/-^ mice fed LFD or HFD. Data are mean ± s.e. P values were determined via one-way ANOVA with Tukey’s multiple comparisons, where #, denotes P<0.05 compared to WT LFD. **i**, Hematoxylin and Eosin (H&E) staining of BAT, SAT, VAT, and liver, as well as oil red-O and Sirius red staining of the liver from female WT and UCP1^-/-^ mice fed LFD or HFD for 16 weeks. **j,k**, RT-qPCR of livers looking at inflammation makers (**j**) and fibrosis markers (**k**) from female WT and UCP1^-/-^ mice fed LFD or HFD for 16 weeks. Data are mean ± s.e. P values were determined via one-way ANOVA with Tukey’s multiple comparisons, where #, denotes P<0.05 compared to WT LFD.

### Both Adipocyte UCP1 and Non-Adipocyte UCP1 Contribute to Body Weight Regulation during DIT

The role of UCP1 in DIT has been studied only in whole-body UCP1 knockout mice, and it remains untested whether adipose-specific loss of UCP1 produces similar effects. Filling this gap is important because UCP1 expression has been reported in non-adipocyte cell populations^18–20^, but whether non-adipocyte UCP1 in these cells has a functional role in body weight, especially during DIT, remains unclear. To dissect the role of adipose tissue-specific UCP1 versus non-adipocyte UCP1 in DIT-associated body weight, adiposity, and liver pathologies, we examined *adiponectin-Cre;Ucp1^floxed^* (*Ucp1^FATKO^*) mice—i.e. mice lacking UCP1 exclusively in adipose tissue—after 16 weeks on either the LFD or HFD at thermoneutrality [Fig. 3a]. Interestingly, unlike whole-body *Ucp1^-/-^* mice on LFD, which exhibited significantly increased total body mass, SAT accumulation, and metabolic efficiency relative to WT [Fig. 1a-1h], *Ucp1^FATKO^* mice on the LFD showed no differences compared to WT [Fig. 3b-3f], suggesting that these phenotypes, while modest over this feeding timespan, are driven by UCP1 loss in non-adipocyte cell populations.

**Figure 3.**
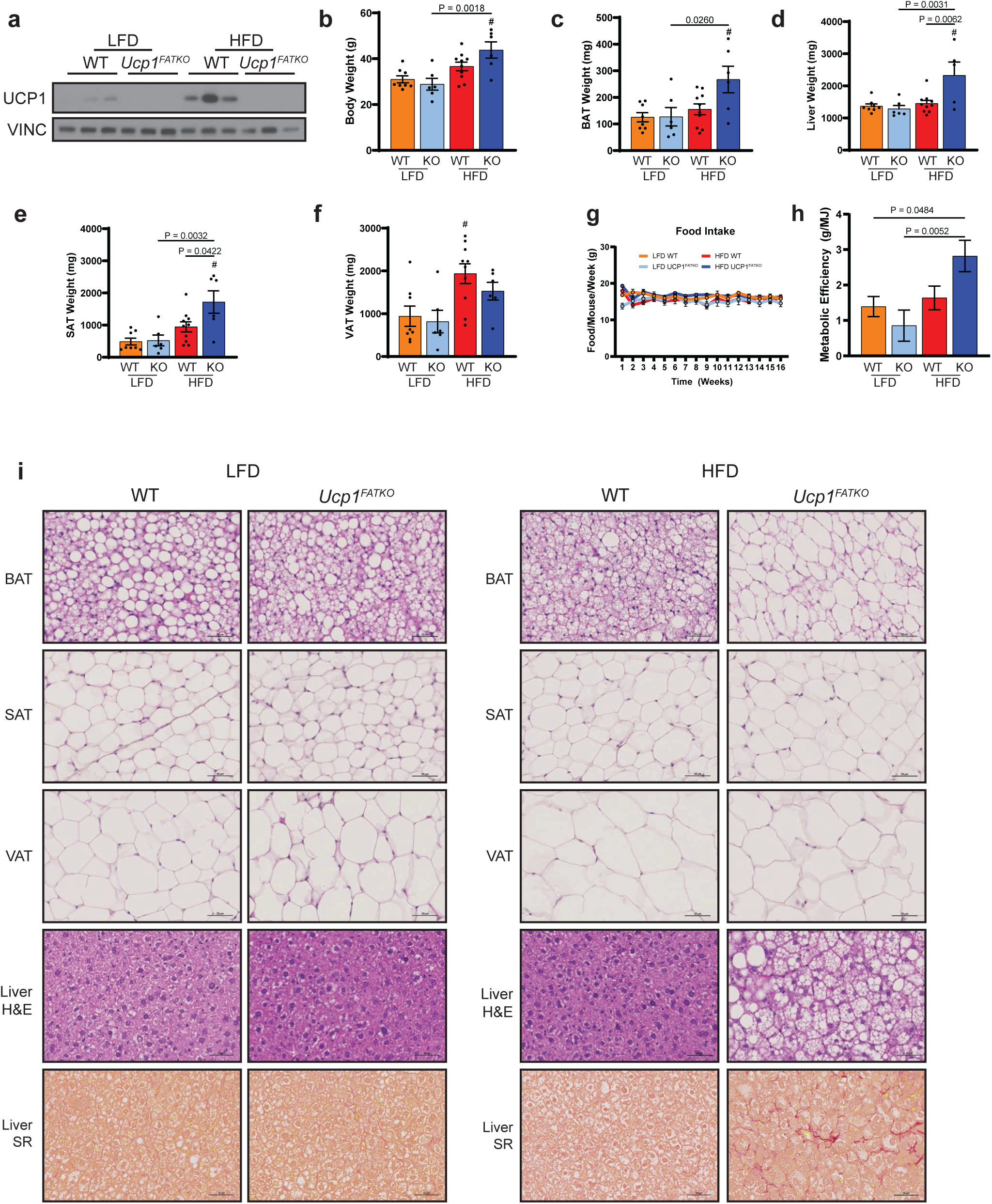
HFD-fed male mice show signs of liver fibrosis in the absence of UCP1 in the adipose tissue. **a**, Immunoblot of BAT from male WT and UCP1^FATKO^ mice fed LFD or HFD for 16 weeks. **b-f**, Body weight (**b**), BAT weight (**c**), Liver weight (**d**), SAT and VAT weights (**e,f**) from male WT and UCP1^FATKO^ mice fed LFD or HFD for 16 weeks. Data are mean ± s.e. P values were determined via one-way ANOVA with Tukey’s multiple comparisons, where #, denotes P<0.05 compared to WT LFD. **g**, Food intake from male WT and UCP1^FATKO^ mice fed LFD or HFD. **h**, Metabolic efficiency calculated by the ratio of body weight gain vs MJ of food consumed from male WT and UCP1^FATKO^ mice fed LFD or HFD. Data are mean ± s.e. P values were determined via one-way ANOVA with Tukey’s multiple comparisons, where #, denotes P<0.05 compared to WT LFD. **i**, Hematoxylin and Eosin (H&E) staining of BAT, SAT, VAT, and liver, as well as Sirius Red staining of the liver from male WT and UCP1^FATKO^ mice fed LFD or HFD for 16 weeks.

In contrast, after HFD feeding, adipose tissue-specific *Ucp1* loss resulted in phenotypic changes paralleling those seen in the whole body *Ucp1^-/-^* mice on HFD. For example, total body mass [Fig. 3b], BAT mass [Fig. 3c], liver mass [Fig. 3d], and SAT mass [Fig. 3e] all increased in *Ucp1^FATKO^* mice on HFD relative to WT LFD mice, while VAT did not [Fig. 3f], just like in *Ucp1^-/-^* mice [Fig. 1b-e]. No difference in food intake was observed between the groups; however, HFD *Ucp1^FATKO^* mice exhibited increased metabolic efficiency compared to LFD-fed mice and a trend toward an increase relative to HFD WT mice [Fig. 3g, 3h]. Consistent with the loss of DIT, the brown adipocytes in the *Ucp1^FATKO^* on HFD lost their multi-locular characteristic [Fig. 3i]. Moreover, the increase in liver mass seen in the HFD *Ucp1^FATKO^* was associated with increased hepatic steatosis and fibrosis [Fig. 3i], again paralleling phenotypes in the whole-body knockout. These findings suggest that while UCP1 in non-adipose tissues may contribute to body weight regulation under the LFD conditions, the protective effects of UCP1 during DIT-induced metabolic stress appear predominantly mediated by adipose tissue-specific UCP1. These data also indicate that the whole-body Ucp1-/- model is valid for investigating the role of brown fat UCP1 in DIT.

### Distinct BAT Gene Signatures Define Diet- and Cold-Induced Thermogenesis

To gain insights into the molecular details of DIT, we performed bulk RNA-sequencing on BAT isolated from WT or whole-body *Ucp1^-/-^* mice on LFD or HFD. To capture early stages of DIT, before more severe metabolic disease arises, we used BAT tissue that was isolated after 8 weeks on HFD, a time point after UCP1 induction was observed in WT mice [Extended Fig. 3a] but preceding the onset of severe hepatic enlargement, inflammation and fibrosis in *Ucp1^-/-^* mice [Extended Fig. 3b-3k].

We first established a DIT gene signature in WT mice by comparing differentially expressed BAT genes between LFD and HFD conditions. This analysis identified 171 upregulated and 152 downregulated genes (>1.5-fold change; FDR < 0.05) [Fig. 4a]. Interestingly, neither *Ucp1* nor *Dio2*, classic markers of CIT, were upregulated at the gene level during DIT [Fig. 4a] (green dots). This suggests that the induction of UCP1 during DIT is through a post-transcriptional mechanism.

**Figure 4.**
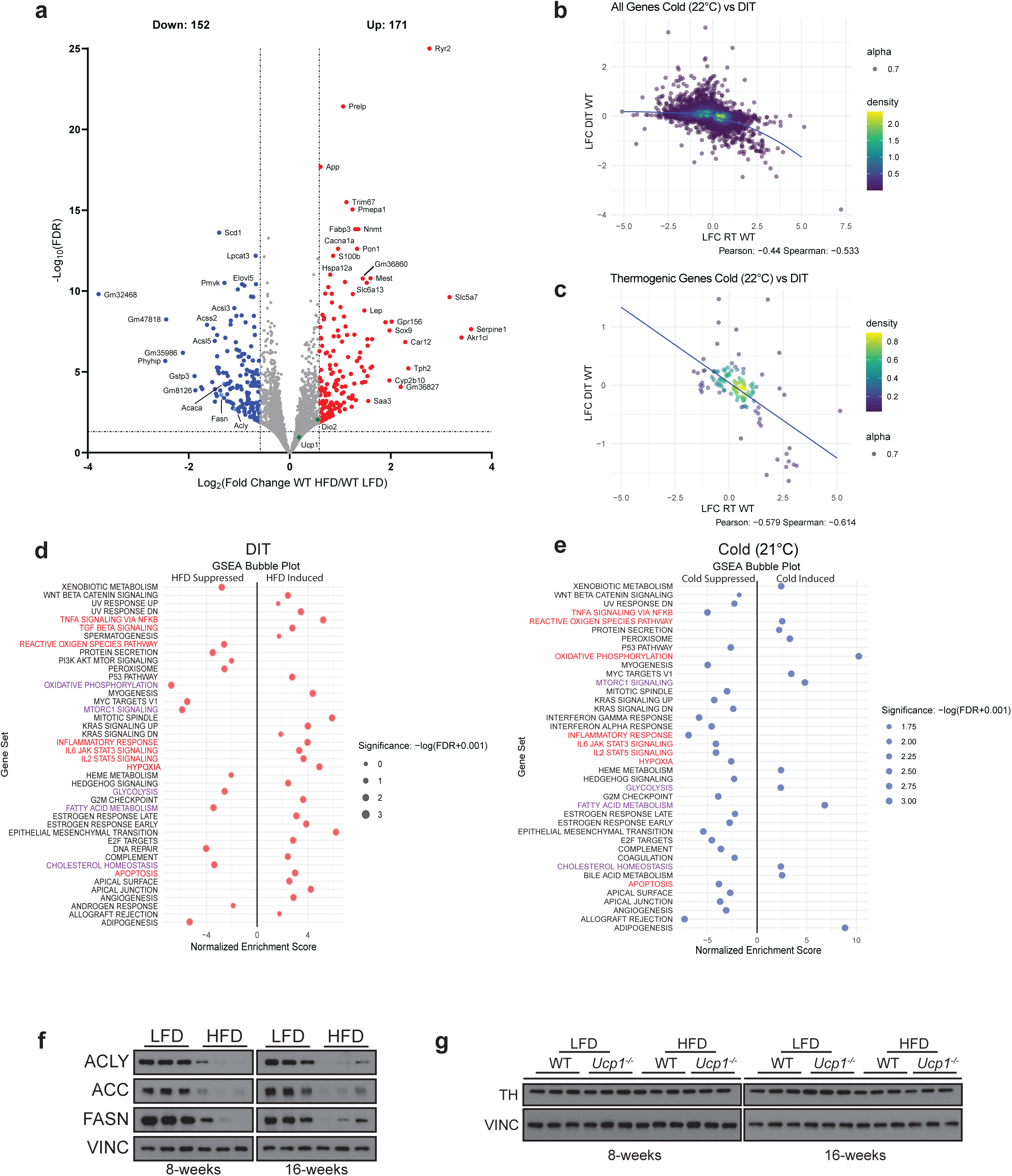
Diet-induced thermogenesis transcriptionally diverges from cold-induced thermogenesis. **a**, Volcano plot depicting genes greater than 1.5-fold and less than 0.05 FDR between WT mice fed a LFD or HFD for 8 weeks. **b**, Correlation between DIT and CIT using all significant genes. **c**, Correlation between DIT and CIT using only significant thermogenic-associated genes. **d**,**e**, Gene set enrichment analysis of BAT comparing DIT (**d**), and CIT (**e**). Red color indicates notable cellular stress and inflammation related signatures and purple indicates notable metabolism related signatures. **f,** Immunoblot of *de novo* lipid synthesis proteins in BAT from male WT and Ucp1^-/-^ mice fed LFD or HFD for 8- or 16 weeks. **g,** Immunoblot showing tyrosine hydroxylase (TH) protein levels in BAT from male WT and Ucp1^-/-^ mice fed LFD or HFD for 8- or 16 weeks.

These findings prompted us to more directly compare DIT gene signatures with our previously generated CIT gene signatures from thermoneutral, cold (22°C), and severe cold (6°C) adapted mice on a standard chow diet^22^. A correlation analysis between DIT and CIT, which compared all genes that were >1.5-fold different (FDR<0.05) in at least one of the gene sets, shows a negative association (Spearman coefficient of -0.533) [Fig. 4b and Extended Fig. 4b]. We also curated a list of 181 genes known to be associated with BAT thermogenesis (see Methods) [Extended Fig. 5]. Of these genes, 125 showed significant changes in expression in either the DIT or CIT gene sets [Extended Fig. 5]. However, correlation analysis yielded a negative Spearman coefficient of -0.614 for cold (22°C) [Fig. 4c] and -0.484 for severe cold (6°C) [Extended Fig. 4c], consistent with DIT and CIT being supported by different transcriptional programs.

The DIT gene set enrichment analysis (GSEA) signature was also strikingly different than that of the CIT GSEA signature [Fig. 4d, 4e and Extended Fig. 4a]. This analysis revealed several pathways induced by DIT, including TNF-α signaling, TGF-β signaling, inflammatory response genes, IL/STAT, hypoxia, and apoptosis signaling [Fig. 4d], which were largely suppressed during CIT at both severe cold (6°C) and cold (21°C) temperatures. One notable exception was hypoxia, which was induced during DIT and suppressed in cold (21°C), but induced in severe cold (6°C) [Extended Fig. 4a]. In contrast, DIT suppresses genes involved in oxidative phosphorylation, mTORC1 signaling, glycolysis, fatty acid metabolism, cholesterol homeostasis, and reactive oxygen species [Fig. 4d], which increase during CIT [Fig. 4e and Extended Fig. 4a]. Moreover, lipid-synthesis genes (*Scd1, Elovl5, Acss2, Acsl3*, and *Acsl5*) were decreased in DIT [Fig. 4a]. Additionally, *de novo* lipogenesis pathway genes (*Acly, Acaca,* and *Fasn*), which are hallmark CIT-associated genes, all were decreased during DIT [Fig. 4a], which we confirmed by immunoblotting [Fig. 4f]. We also noted that tyrosine hydroxylase (TH), a reporter used to indicate sympathetic nervous system stimulation of BAT during CIT^5^, was unaffected by either 8 or 16 weeks of HFD despite UCP1 induction [Fig. 4g]. Taken together, these data suggest that divergent regulatory programs drive DIT and CIT.

### UCP1 Suppresses Inflammation and Stress Pathways in BAT During DIT

We next examined UCP1’s role in regulating BAT gene expression during DIT. In LFD-fed mice, *Ucp1* loss resulted in 656 upregulated and 117 downregulated genes of which *Ucp1* was the most decreased gene [Fig. 5a]. During DIT, *Ucp1* loss resulted in much more profound effects, with 1106 upregulated and 611 downregulated genes [Fig. 5b]. GSEA indicated oxidative phosphorylation, mTOR, glycolysis, fatty acid metabolism, and bile acid metabolism genes decreased with *Ucp1* loss on LFD, while cholesterol synthesis increased [Fig. 5c]. Many inflammatory and stress signatures (inflammatory response, TNFA, JAK/STAT, apoptosis, hypoxia) were further increased by the loss of *Ucp1*, suggesting that UCP1 basally suppresses these stress responses even on the LFD condition [Fig. 5c]. During HFD conditions, pathway analysis also showed a decrease in the metabolic signatures of oxidative phosphorylation and fatty acid metabolism [Fig. 5d]. However, unlike LFD, glycolysis and mTOR signature genes increased in the HFD condition [Fig. 5d]. *Ucp1* loss also increased a broad spectrum of inflammatory and stress gene signatures in the HFD state, including TNFA signaling, JAK/STAT, apoptosis, hypoxia, interferon-gamma and -alpha responses, and reactive oxygen species genes [Fig. 5d]. We noted that *Ucp2* was highly upregulated regardless of diet [Fig. 5a,5b] possibly as a compensatory mechanism^23–25^. These data further indicate UCP1 functioning during DIT to reduce overall metabolic stress and inflammation.

**Figure 5.**
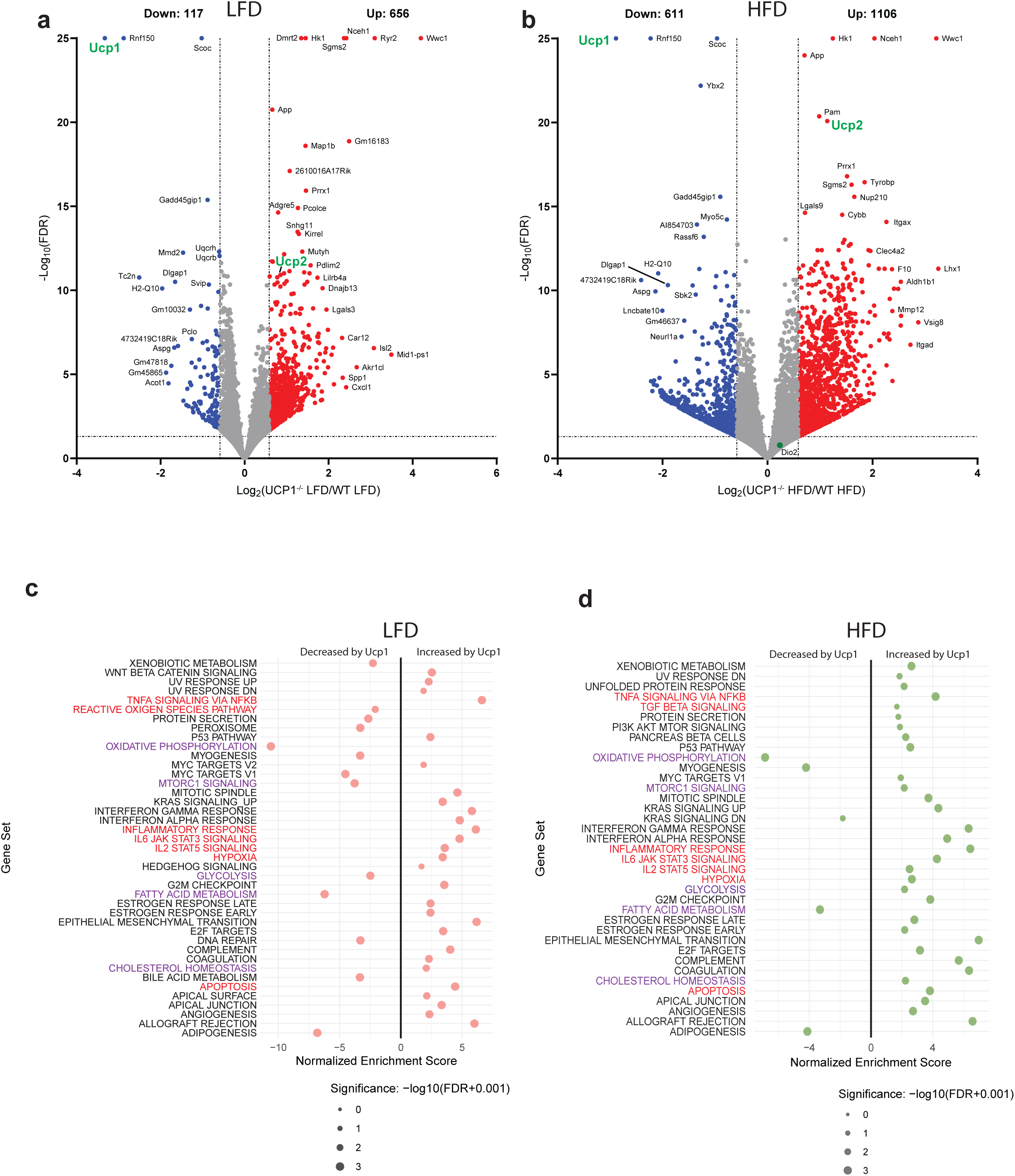
UCP1 loss reveals altered metabolic and transcriptional profiles. **a**,**b**, Volcano plot showing differential gene expression in BAT between UCP1^-/-^ LFD and WT LFD mice (**a**), and UCP1^-/-^ HFD and WT HFD mice (**b**). **c**,**d**, Gene set enrichment analysis in BAT comparing loss of UCP1 on LFD (**c**), and HFD (**d**). Red color indicates notable cellular stress and inflammation related signatures and purple indicates notable metabolism related signatures.

### DIT Augments BAT Uptake and Release of Circulating Nutrients

Having established that DIT differs markedly from CIT at the level of gene expression, we next asked whether these two thermogenic states also differ in their use of circulating nutrients. Activated BAT is proposed to function as a catabolic sink, which may contribute to its systemic metabolic benefits^15^. To explore this concept during DIT, we investigated how DIT alters BAT consumption of circulating nutrients in a UCP1-dependent manner by leveraging BAT-specific arteriovenous (AV) metabolomics [Fig. 6a] ^26,27^. We performed this assay at the same time point when we examined gene profiles using RNA-sequencing [Extended Fig. 4], after 8 weeks of LFD/HFD feeding.

**Figure 6.**
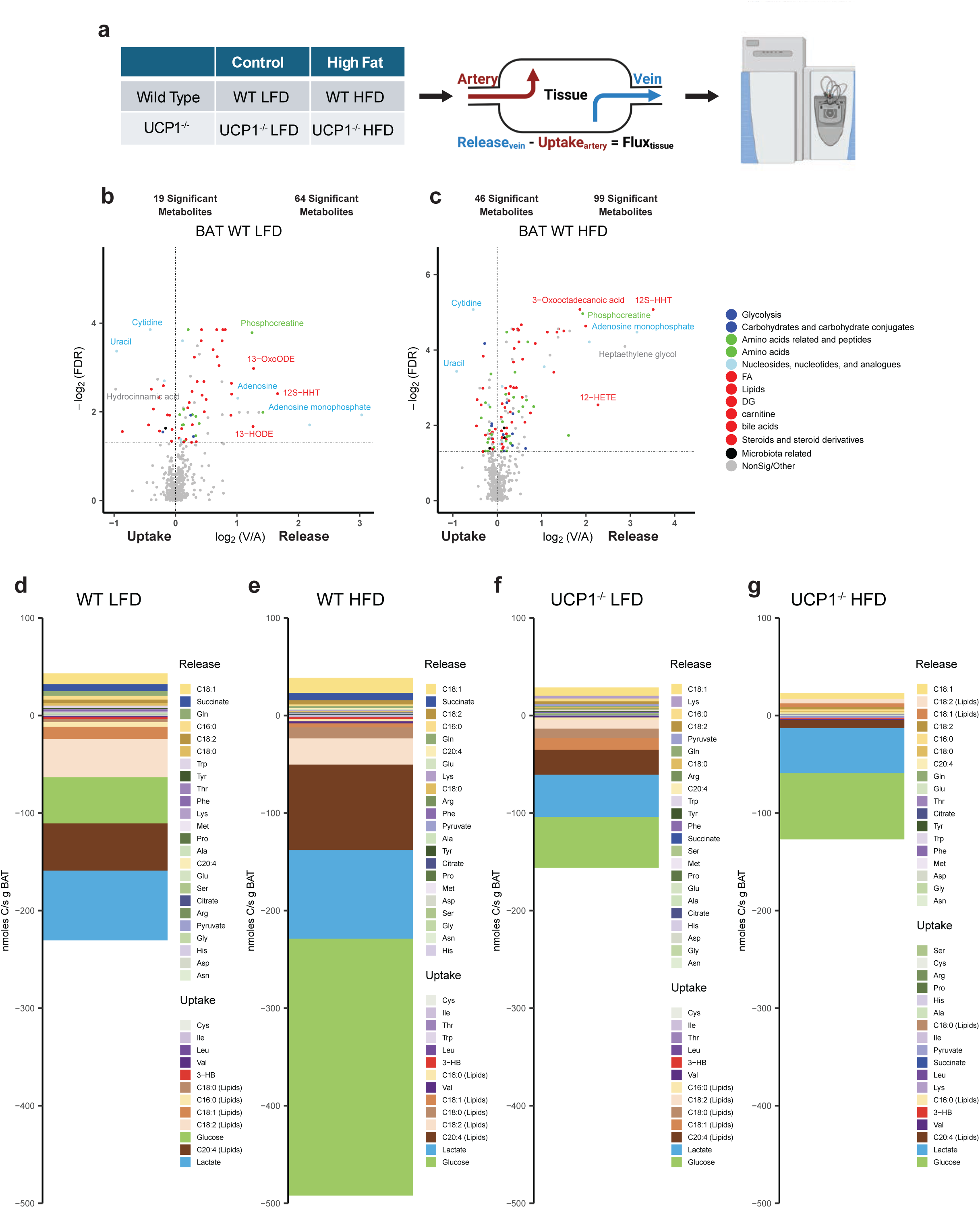
Arteriovenous metabolomics reveals altered nutrient utilization during diet-induced thermogenesis. **a**, Schematic detailing AV metabolomics workflow. **b**, Untargeted metabolite profiling showing release and uptake in WT LFD BAT. **c**, Untargeted metabolite profiling showing release and uptake in WT HFD BAT. **d-g**, Quantitative carbon flux measurements in all four conditions WT LFD (**d**), WT HFD (**e**), UCP1^-/-^ LFD (**f**), UCP1^-/-^ HFD (**g**). Metabolites are ordered based on their relative contributions from greatest to least.

The resulting data revealed that BAT consistently takes up major circulating nutrients such as glucose, lactate, and ketones while it releases fatty acids regardless of whether the mice are consuming LFD or HFD [Fig. 6b,6c and Extended Fig. 6a-6g and Extended Data Table 1]. However, untargeted metabolomics displayed that DIT markedly increased the number of metabolites consumed by BAT (from 64 to 99) and released by BAT (from 19 to 46), reflecting augmented metabolic activity [Fig. 6b,6c and Extended Data Table 1]. Intriguingly, when we performed similar AV analysis across the liver and small intestine during DIT to examine potential interactions between BAT and these metabolically active organs, we found no profound impact of DIT on the liver and intestine metabolite uptake or release [Extended Fig. 6a-6l; Extended Fig. 7a-b,7e-f and Extended Data Table 2 and 3]. Thus, while DIT is associated with stimulated BAT metabolite uptake and release, it appears to exert a minimal effect on hepatic or intestinal metabolite tissue flux.

### Glucose is the Major Carbon Source for BAT during DIT

To quantitatively evaluate the impact of DIT on BAT metabolite uptake and release, focusing on primary building blocks and energy sources, we calculated the net influx and efflux of 36 highly abundant circulating metabolites across BAT^26,28^. Briefly, to calculate the total carbon flux of BAT (Fc):

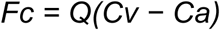

where Q is BAT blood flow rate (Extended Fig. 7c), as measured with color and pulsed-wave Doppler ultrasound. Cv and Ca are blood concentrations of carbon carried by each metabolite in the Sulzer’s venous and arterial blood. This quantitative analysis revealed a 2-fold increase in total carbon consumption by BAT during DIT (492.1 nmol C s^−1^g^−1^ BAT) compared to control (230.4 nmol C s^−1^g^−1^ BAT) (see Methods). This was mostly attributed to a ∼5-fold increase in glucose uptake [Fig. 6d,6e] (green bar) from 47.4 (LFD conditions) to 263.2 nmol C s^−1^g^−1^ BAT weight (HFD conditions). Consequently, BAT during DIT obtained >50% of total carbon from glucose, whereas BAT in control condition obtained ∼25% [Fig. 6d,6e]. In contrast, circulating fatty acids (C20:4, C18:2, C18:1, C18:0, C16:0) contributed less (∼27%) to BAT during DIT compared to the control condition (∼47%) [Fig. 6d,6e], indicating a fuel utilization shift in the BAT away from using lipids towards increased glucose utilization during DIT, which was surprising given the high-lipid content of the HFD.

We also performed a similar quantitative analysis in the UCP1-deficient mice during DIT. Strikingly, loss of UCP1 almost completely blunts HFD-induced glucose uptake by BAT while minimally impacting other metabolites [Fig. 6f,6g and Extended Fig. 8a,8b and Extended Data Table 1]. UCP1 loss also barely affected metabolite uptake and release by the liver or small intestine, with only a modest blunting of released metabolites with UCP1 loss on LFD [Extended Fig. 7c-d,7g-h and Extended Data Table 2 and 3]. Thus, despite the high fat content of the diet that stimulates DIT, glucose is the primary circulating fuel consumed by BAT during this process. This suggests a specific link between UCP1 and glucose utilization during DIT that is independent of sympathetic stimulation and the conventional thermogenic gene expression program induced by cold.

## Discussion

The regulation and function of BAT metabolic activity during DIT are poorly understood, largely because most prior studies have mainly focused on BAT responses to cold exposure rather than nutrient stimulation. Moreover, investigations of DIT have primarily focused on the contribution of whole-body UCP1 to weight gain and adiposity^5,7,8,10,16^, leaving the tissue-specific role of UCP1 and its broader impact on whole-body metabolism, largely unexplored. It has also been unclear whether DIT and CIT employ biologically similar pathways for thermogenic activities, and whether male and female BAT responds similarly to nutrient stress at thermoneutrality. Here, we fill these gaps, advancing a new view of how nutrients can stimulate brown fat activity independently of cold exposure.

By comprehensively characterizing the physiological and molecular impact of diet-induced UCP1 upregulation in BAT, we reveal an unappreciated impact of DIT on liver metabolic disease. We also uncovered a potential role for non-adipocyte UCP1 in regulating body weight. Several studies have suggested the presence of UCP1 outside of adipose tissues^18–20^. Although UCP1 expression in non-adipose tissues is far lower than in brown or beige fat, its cumulative abundance across these tissues could still influence body weight regulation, as suggested by this study, and warrants further investigation in both mice and humans.

We also found that DIT substantially diverges from CIT at the transcriptome level, with notable differences including the increase in inflammatory and stress responses associated with DIT but not with CIT, and the lack of increased fatty acid synthesis, a hallmark of cold-activated BAT^22,29^. Interestingly, however, BAT-specific arteriovenous metabolomics during DIT revealed that BAT prefers glucose as a fuel source during DIT, which is similar to BAT in cold-adapted mice with free access to a standard diet^26^. This finding was unexpected, given the high lipid content of the HFD and the observation that CIT and DIT exhibit such different transcriptional profiles, as suggested by their strong negative correlation. The notable commonality between DIT and CIT is the induction of UCP1 protein levels, highlighting a functional link between UCP1 and glucose utilization. This link could reflect an increased demand for glycolytic ATP production during uncoupling respiration, cytosolic production of antioxidants, or some other aspect of UCP1-dependent metabolism that is best supported by glucose.

Nevertheless, while increased UCP1 levels and glucose utilization are shared between diet- and cold-induced thermogenesis, these modes of BAT activation appear to be fundamentally different processes. One possibility suggested by our data is that DIT is an adaptive response to nutrient stress, which would be consistent with UCP1’s known role in mitigating mitochondrial stress^23–25,30^. In contrast, cold activates a highly specialized BAT metabolic program that evolved specifically to defend body temperature under sub-thermoneutral conditions, potentially building upon ancestral roles of uncouplers in protecting against metabolic stress.

Given the prevalence of thermoneutral environments and high-fat diets in modern society, our findings provide key insights into the molecular mechanisms, contributing organs, and sex-specific differences in diet-induced thermogenesis, with potential translational relevance for targeting metabolic disease.

## Acknowledgments

We thank Lawrence Kazak for providing the UCP1 floxed mice. We thank Benjamin Clayton, Alex Boucher Jr., and Emma Mohlmann for their assistance with animal husbandry and discussion. We thank all members of the Jang and Guertin laboratories for the discussion.

## Author contributions

D.A.G., C.J., and J.A.H. conceptualized the study. J.A.H. performed almost all animal experiments, sample collection, and sample preparation. J.L. and C.J. performed all the serum sample preparation and analysis for the arteriovenous metabolomics experiments. R.L. and L.J.Z. performed RNAseq analysis. E.D.K., F.D.S., M.G. all assisted with dissections, sample prep, and animal husbandry. C.A.L. assisted with metabolic analysis. T.P.F. performed Doppler imaging and analysis. N.M. and J.E.H. performed and analyzed flow cytometry for immune cells. Q.C. and S.M.F. assisted with animal dissections and animal husbandry. H.L. performed animal genotyping and assisted with animal colony management. A.C.M. assisted with liver fibrosis experimentation and interpretation. J.A.H., D.A.G., and C.J. wrote the manuscript.

## Data and materials availability

All data needed to evaluate the conclusions in the paper are present in the paper

## Methods

### Mice

All research complies with all relevant ethical regulations set by the NIH, IACUC, and IBC. C57BL/6J mice stocks were purchased from Jackson Laboratories and maintained in the lab. UCP1^-/-^ mice were obtained from Jackson Labs and created as described in^31^. Mice were bred and housed at 22°C with a 12h light/12h dark cycle, with free access to normal water and a standard chow diet (Prolab Isopro® RMH 3000). All animal experiments were approved by the University of Massachusetts Chan Medical School Institutional Animal Care and Use Committee. For all diet experiments, 6-week-old mice were placed in a temperature-controlled room set to 30°C. Mice were adapted at 30°C for four weeks with free access to food and acid water and maintained on the standard day/night light cycle. At 10-weeks old, mice were placed on a 45% high-fat diet (ResearchDiet#D12451i) or calorically matched control diet (ResearchDiet#D12450Hi) for 8- or 16-weeks. All mice had free access to food and acid water and were maintained on the standard day/night light cycle.

### AV blood sampling and tissue processing

*Ad libitum* fed mice were anesthetized with isoflurane using the drop method and placed on a dissection tray with a nose cone. Blood collection was completed within 5 minutes of anesthesia administration. An incision below the shoulder was made and the interscapular brown adipose was lifted toward the head to expose the intact Sulzer’s vein. Fine surgical scissors were used to cut the Sulzer’s vein and ∼40 μL of blood was collected with capillary tubes without EDTA or heparin coating. Next, the mouse was flipped and cut open to expose the diaphragm. The left ventricle was then punctured through the diaphragm and ∼50 μL blood was collected with a 29G, ½ mL insulin syringe. Next, the intestines were moved to reveal the portal vein. A 29G, ½ mL insulin syringe was used to puncture the portal vein. Once punctured, ∼40 μL of blood was collected. Finally, the liver upper lobes of the liver were pulled down toward the tail to reveal the hepatic vein, which was punctured using a 29G, ½ mL insulin syringe and ∼40 uL of blood was collected. For serum collection, blood samples were placed on ice in an anticoagulant-free tube for 20 min, followed by centrifugation at 10,000 x g for 10 minutes at 4°C. The resulting supernatant was stored at -80°C and analyzed within a week. The interscapular BAT depots and liver tissue were snap-frozen using a liquid nitrogen-cooled Wollenburg clamp ^32^. Tissues were put into 2 mL Eppendorf tubes with a pre-cooled 5 mm metal bead. The CryoMill (Retsch, Newtown, PA) was pre-cooled before loading the 2 mL tubes. Once pre-cooled, the tubes were loaded, and all tissues were milled at 25 Hz for 0.5-2 minutes. Metal beads were removed, and tissue powder was weighed out into dry ice pre-cooled 1.5 mL Eppendorf tubes for metabolite extraction, immunoblot, or gene expression analysis.

### Tissue histology

Tissue pieces were fixed in 10% formalin or embedded in OCT and frozen for oil red-O staining. Embedding, sectioning, Hematoxylin and Eosin (H&E) staining, Oil red-O and picrosirius red staining was done by the UMCMS Morphological Core facility. Full brightfield slides scans were taken at 20x using a Zeiss Axio-Scan.Z1. Zeiss Zen Blue was used for image processing.

### Glucose tolerance tests

Mice after either 6-weeks of LFD/HFD or 14-weeks of LFD/HFD were used. Mice were fasted for 14-hours prior to glucose tolerance test, and all tests were started between 8-9am. Glucose was injected via IP injection at a dose of 1g/kg in phosphate-buffered saline, unless otherwise specified. Tail blood was measured on a glucometer (GE-100) at 0, 15, 30, 60, and 120 minutes after glucose administration.

### Soluble metabolite measurements using LC-MS

For serum metabolite extraction, 5 μL of serum was mixed with 150 μL 40:40:20 of acetonitrile:methanol:water mixture, vortexed, and centrifuged at 16,000 x g for 10 min at 4°C. 3 μL of supernatant was injected to LC-MS. A quadrupole orbitrap mass spectrometer (Q Exactive; ThermoFisher Scientific) operating in negative or positive ion mode was coupled to a Vanquish UHPLC system (ThermoFisher Scientific) with electrospray ionization and used to scan from m/z 70 to 1,000 at 2 Hz, with a 140,000 resolution. LC separation was achieved on an XBridge BEH Amide column (2.1 x 150 mm^2^, 2.5μm particle size, 130 Å pore size; Waters Corporation) using a gradient of solvent A (95:5 water: acetonitrile with 20 mM of ammonium acetate and 20 mM of ammonium hydroxide, pH 9.45) and solvent B (acetonitrile). Flow rate was 150μl/min. The LC gradient was: 0 min, 85% B; 2 min, 85% B; 3 min, 80% B; 5 min, 80% B; 6 min, 75% B; 7 min, 75% B; 8 min, 70% B; 9 min, 70% B; 10 min, 50% B; 12 min, 50% B; 13 min, 25% B; 16 min, 25% B; 18 min, 0% B; 23 min, 0% B; 24 min, 85% B; and 30 min, 85% B. The autosampler temperature was 5°C. Metabolite concentrations were determined by authentic synthesized standards from Sigma. Data were analyzed using the MAVEN software (build 682, http://maven.princeton.edu/index.php) and Compound Discoverer software (Thermofisher Scientific).

### Echocardiography and Vascular Imaging

All imaging was done in the UMass Chan Medical School Cardiovascular & Surgical Models Core (IACUC protocol #20220006) and as previously described in^26^. The back and chest were treated with depilatory cream. Animals were induced with 2.0% isoflurane mixed with 0.5 L/min 100% 0_2_ and gently affixed in the prone position to the heated physiologic platform of the Vevo 3100 imaging system (Visualsonics, Toronto, ON, Canada). Electrode cream was applied to each limb. Body temperature was continually monitored and maintained at ∼37°C with a rectal temperature probe. Isoflurane was administered by nose cone and the concentration reduced to 1.0 % isoflurane mixed with 0.5 L/min 100% 0_2_. All animals were imaged at a heart rate of at least 400-500 beats per minute (bpm), any animals below 400 bpm were excluded from the analysis. A 50 MHz transducer (MX550S) was used to find the Sulzer vein. Vein flow was verified with color flow doppler. Pulsed wave doppler was used to measure flow in the Sulzer’s vein. The velocity time integral (VTI) of flow was measured for three consecutive beats at end expiration. The diameter of the vein was measured with 2D doppler using the leading edge to leading edge technique. Flow in the Sulzer vein was calculated by multiplying the average VTI of vein flow by the cross-sectional area of the vein (*CAS* = *πr*^2^) and heart rate (bpm). Flow in the Sulzer’s vein was normalized to the median weight of BAT tissue across all studies. Mice were then placed supine for echocardiography. 2D and M-mode images were obtained in the parasternal long and parasternal short axis as previously described ^33,34^. Image analysis was performed off-line using Vevo Lab image analysis software. LV volumes were derived from m-mode measurements using the following formulas: LV diastolic volume ((7.0/(2.4 + LVID;d))*LVID;d^3^ and LV systolic volume ((7.0/(2.4 + LVID;s))*LVID;s^3^. Cardiac output was calculated by the following equation (CO=stroke volume x heart rate). LV mass was calculated using the following formula (1.053*((LVID;d + LVPW;d + IVS;d) ^3^-LVID;d^3^).

### Immunoblot analysis

For immunoblot analysis of surgically dissected tissue depots, tissues were cryomilled and lysed in RIPA buffer (150 mM NaCl, 50 mM HEPES at pH 7.4, 0.1% SDS, 1% Triton X-100, 1% glycerol, 2 mM EDTA, 0.5% deoxycholate) containing a protease and phosphatase inhibitor cocktail. Protein lysates were mixed with 5X SDS sample buffer and boiled, separated by SDS-PAGE, transferred to polyvinylidene difluoride (PVDF) membrane, and subjected to immunoblot analysis.

### qPCR analysis

RNA was isolated from tissues using Qiazol (QIAGEN) and RNeasy kit (QIAGEN). 1 μg of RNA was reverse transcribed to cDNA using a high-capacity cDNA reverse transcription kit (#4368813, Applied Biosystems). Quantitative RT-PCR (qRT-PCR) was performed in 10uL reactions using a StepOnePlus real-time PCR machine from Applied Biosystems using SYBR Green PCR master mix (CWBio#CW0955) according to manufacturer instructions. Melting curves were run on every plate for all genes to ensure the efficiency and specificity of the reaction. TATA-box binding protein (Tbp) gene expression was used for normalization. Data acquisition was performed with Applied Biosystems StepOne Software.

### Gene Expression Analysis

TN, RT, and SCRNA sequencing data are from^22^. Samples for DIT RNA sequencing, RNA was isolated from tissues using Qiazol (QIAGEN) and an RNeasy kit (QIAGEN), and sequencing was performed by Azenta NGS. RNASeq analysis was performed with OneStopRNAseq^35^. Paired-end reads were aligned to mouse genome mm10, with star_2.5.3a^36^, annotated with GENCODE GRCm38.p6 annotation release 25^37^. Aligned exon fragments with mapping quality higher than 20 were counted toward gene expression with featureCounts_1.5.2^38^. Differential expression (DE) analysis was performed with DESeq2_1.20.0^39^. Within DE analysis, ’ashr’ was used to create log2 Fold Change (LFC) shrinkage for each comparison^40^. Expression heatmaps were created with pheatmap^41^, zscore of TPM were used for visualization. Gene set enrichment analysis were performed with GSEA^42^. Mouse Molecular Signatures Database (MSigDB)^43^ was used as the gene sets database. The multi-bubble plot for GSEA results only included enriched gene sets with FDR < 0.05.

### Flow cytometry

Single cell suspension was done from BAT and liver using gentleMACS^TM^ tissue dissociator (Miltenyi Biotec, Auburn, CA) using collagenase A (Roche, Indianapolis, IN) and IV (Sigma-Aldrich, St. Louis, MO), respectively. Isolated cells were stained with Zombie Aqua Viability Kit and for CD45 Bv570 (30-F11), CD19 A488 (6D5), CD3 APC-Cy7 (17A2), NK1.1 Bv650 (PK136), CD11c Bv785 (N418), CD11b PercP (M1/70), F4/80 PE/Dazzle 594 (BM8), Ly6C A700 (HK1.4), Ly6G Bv605 (1A8), MHC-II PE (M5/114.15.2), Tim-4 PE-Cy7 (RMT4-54), ESAM APC (1G8/ESAM), CD206 Bv421 (C068C2) and CD64 APC (X54-5/7.1) (Biolegend, San Diego, CA). All antibodies were used in a 1:200 dilution. Antibodies were validated by Biolegend in mouse splenocytes and extensive literature has been published using these antibodies. Data was acquired on a Cytek^®^ Autora (Cytek, Freemont, CA) and was analyzed with FlowJo v10.9.0. The gating strategy is detailed in Fig. 2a.

### Olink proteomics

Plasma obtained from the portal vein was used for the proximity extension assay (PEA) from Olink^®^ Proteomics. The Olink Target 48 Mouse Cytokine assay was performed, and results were calculated as pg/ml or normalized protein expression (NPX) using Olink^®^ NPX signature software v1.14.0.

### Statistics and Reproducibility

Metabolic heatmaps were generated using Metaboanalyst and R 4.1.1 software (gplots). Statistical analysis was performed using Graphpad Prism 9.0 and R software (rstatix). When two groups were compared, a two-tailed, unpaired Student’s t-test was used to calculate P values, with P < 0.05 used to determine statistical significance. FDR correction conducted with Benjamini-Hochberg. When >2 groups were compared, a one-way ANOVA was employed. Tukey’s method was used to correct for multiple comparisons. All immunoblots in this study are representative of at least three independent experiments.

## Extended Figure Legends

**Extended Figure 1. Characterization of DIT in male WT mice at 8- and 16-weeks. a,** Immunoblot showing UCP1 protein expression in BAT of 8- and 16-week LFD or HFD fed male WT mice. **b**, Hematoxylin and Eosin (H&E) staining of BAT from 8- and 16- week LFD or HFD fed male WT mice. **c,d,** Body weight (**c**) and tissue weights (**d**) from 8-week LFD or HFD fed male WT mice. Data are mean ± s.e. P values were determined via Students T-test. **e**, Hematoxylin and Eosin (H&E) staining of SAT, VAT, and Liver, as well as oil red-O of liver from 8-week LFD or HFD fed male WT mice. **f**, Area under the curve of IPGTT of LFD or HFD fed male WT mice. **g,h,** Body weight (**g**) and tissue weights (**h**) from 16-week LFD or HFD fed male WT mice. Data are mean ± s.e. P values were determined via Students T-test. **i**, Hematoxylin and Eosin (H&E) staining of SAT, VAT, and Liver, as well as oil red-O of liver from 8-week LFD or HFD fed male WT mice. **j**, Area under the curve of IPGTT of LFD or HFD fed male WT mice.

**Extended Figure 2. *Ucp1^-/-^* HFD male mice have increased macrophage infiltration in the liver and circulating inflammatory markers. a**, Gating for all flow cytometry experiments. **b-i,** Flow cytometry showing immune cell proportion and numbers in the livers of 16-week LFD or HFD fed male WT or *Ucp1^-/-^* mice. Data are mean ± s.e. P values were determined via one-way ANOVA with Tukey’s multiple comparisons, where #, denotes P<0.05 compared to WT LFD. **j-l,** Olink showing circulating inflammatory markers in the portal vein of 16-week LFD or HFD fed male WT or *Ucp1^-/-^* mice. Data are mean ± s.e. P values were determined via one-way ANOVA with Tukey’s multiple comparisons.

**Extended Figure 3. HFD-fed male mice do not develop liver fibrosis in the absence of UCP1 at 8 weeks. a**, Immunoblot of BAT from male WT and UCP1^-/-^ mice fed LFD or HFD for 8 weeks. **b-f**, Body weight (**b**), BAT weight (**c**), Liver weight (**d**), SAT and VAT weights (**e,f**) from male WT and UCP1^-/-^ mice fed LFD or HFD for 8 weeks. Data are mean ± s.e. P values were determined via one-way ANOVA with Tukey’s multiple comparisons, where #, denotes P<0.05 compared to WT LFD. **g**, Food intake from male WT and UCP1^-/-^ mice fed LFD or HFD. **h**, Metabolic efficiency calculated by the ratio of body weight gain vs MJ of food consumed from male WT and UCP1^-/-^ mice fed LFD or HFD. Data are mean ± s.e. P values were determined via one-way ANOVA with Tukey’s multiple comparisons, where #, denotes P<0.05 compared to WT LFD. **i**, Hematoxylin and Eosin (H&E) staining of BAT, SAT, VAT, and liver, as well as oil red-O and Sirius red staining of the liver from male WT and UCP1^-/-^ mice fed LFD or HFD for 8 weeks. **j,k**, RT-qPCR of livers looking at inflammation makers (**j**) and fibrosis markers (**k**) from male WT and UCP1^-/-^ mice fed LFD or HFD for 8 weeks. Data are mean ± s.e. P values were determined via one-way ANOVA with Tukey’s multiple comparisons, where #, denotes P<0.05 compared to WT LFD.

**Extended Figure 4. Comparative gene set enrichment analysis reveals a divergence between DIT and CIT. a,** Gene set enrichment analysis of severe cold (SC). Red color indicates notable cellular stress and inflammation related signatures and purple indicates notable metabolism related signatures. **b,** Correlation plot of all genes comparing severe cold and DIT. **c,** Correlation plot of thermogenic-associated genes comparing severe cold and DIT.

**Extended Figure 5. Thermogenic-associated genes in DIT and CIT. a,** Heatmap showing thermogenic-associated genes in WT room temperature (22°C), WT thermoneutral (30°C), WT LFD, and WT HFD male mice.

**Extended Figure 6. Arteriovenous metabolomics of BAT, liver, and small intestines during DIT. a-l**, Bar graphs of key released and consumed fuels across all three tissues from WT and UCP1^-/-^ LFD and HFD fed male mice at 8 weeks. Data are mean ± s.e. #, P values were compared to null value (zero exchange) by one-tailed one-sample t-test. P values for group differences were determined via one-tailed ANOVA with Tukey’s honestly significant difference (HSD).

**Extended Figure 7. Arteriovenous metabolomics of the liver and small intestines during DIT. a-d**, Volcano plots of untargeted metabolite profiles from WT LFD livers (**a**), WT HFD livers (**b**), UCP1^-/-^ LFD liver (**c**), and UCP1^-/-^ HFD liver (**d**). **e-h**, Volcano plots of untargeted metabolite profiles from WT LFD small intestines (**e**), HFD small intestines (**f**), UCP1^-/-^ LFD small intestines (**g**), and UCP1^-/-^ HFD small intestines (**h**).

**Extended Figure 8. Arteriovenous metabolomics of BAT with UCP1 loss. a,b,** Volcano plots of untargeted metabolite profiles from UCP1^-/-^ LFD BAT (**a**), and UCP1^-/-^ HFD BAT (**b**). **c**, Blood flow values in BAT from WT and UCP1^KO^ LFD and HFD fed male mice at 8 weeks. Data are median values.

## Extended Data Tables

**Extended Data Table 1. List of metabolites significantly taken up or released by BAT.**

**Extended Data Table 2. List of metabolites significantly taken up or released by Liver.**

**Extended Data Table 3. List of metabolites significantly taken up or released by Small Intestine.**

