## Extended Data for "UCP1 Mitigates Hepatic Steatosis and Fibrosis Independent of Cold Exposure"

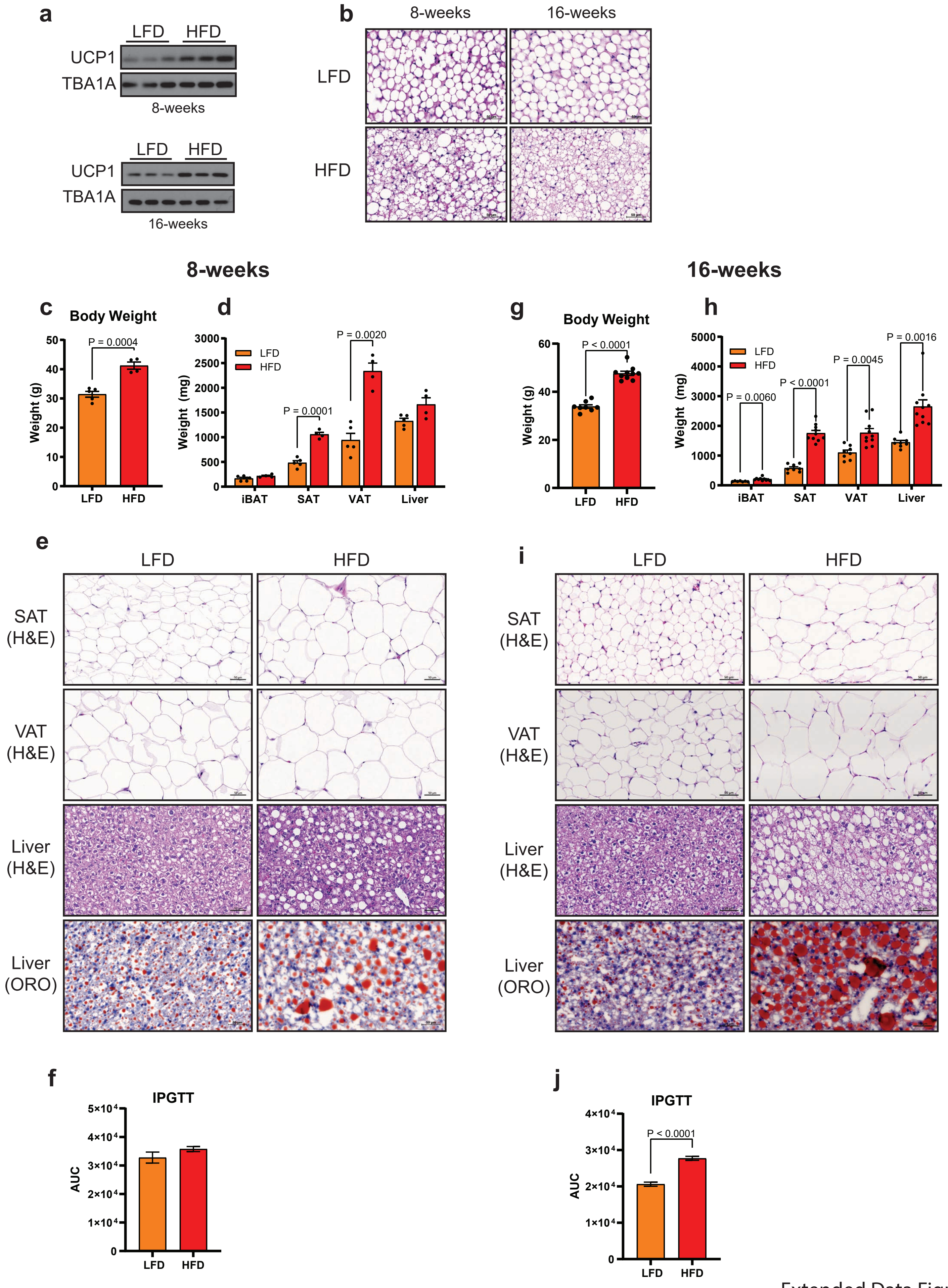


Extended Data Figure 1
Supports Figure 1

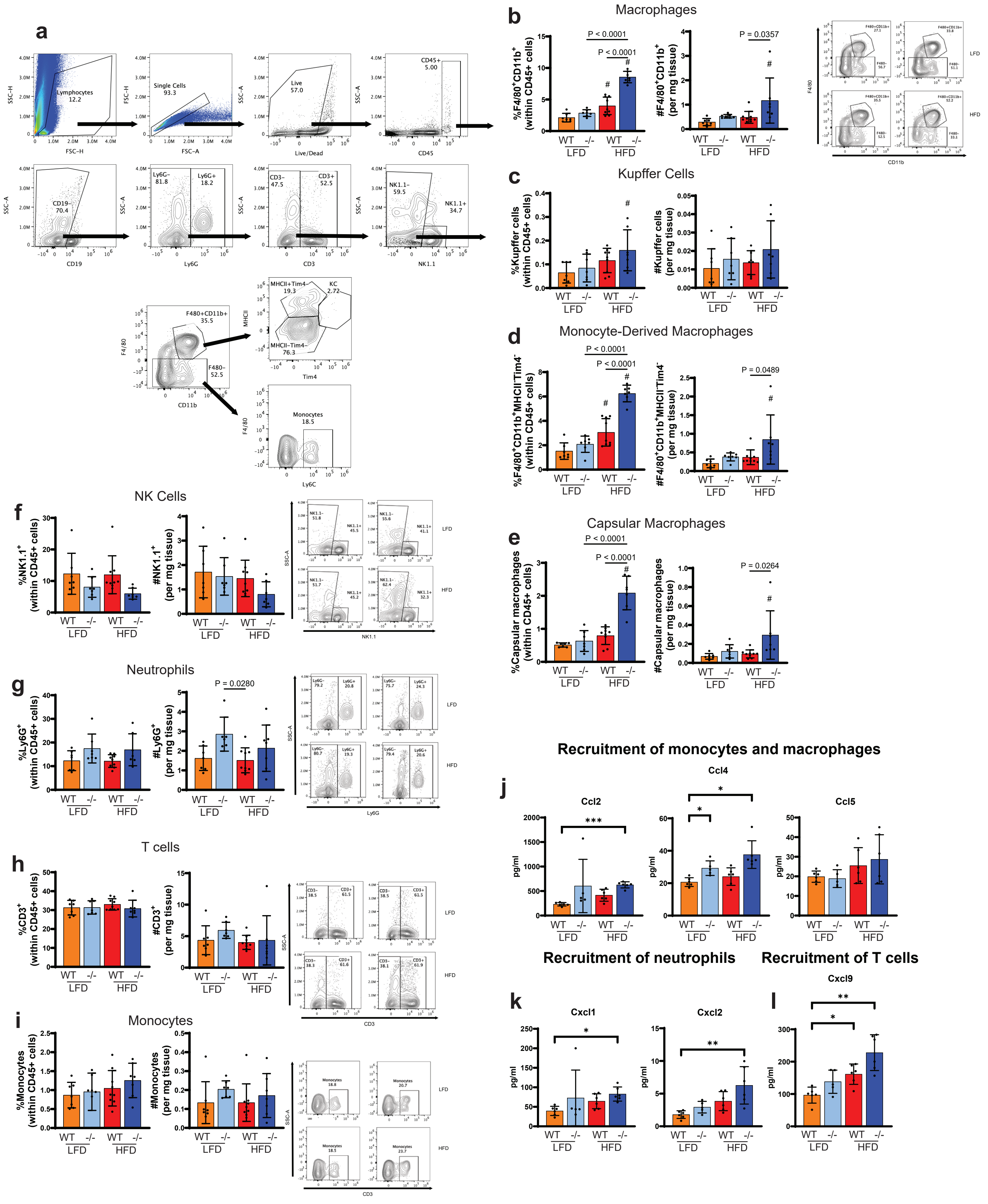


Extended Data Figure 2
Supports Figure 1

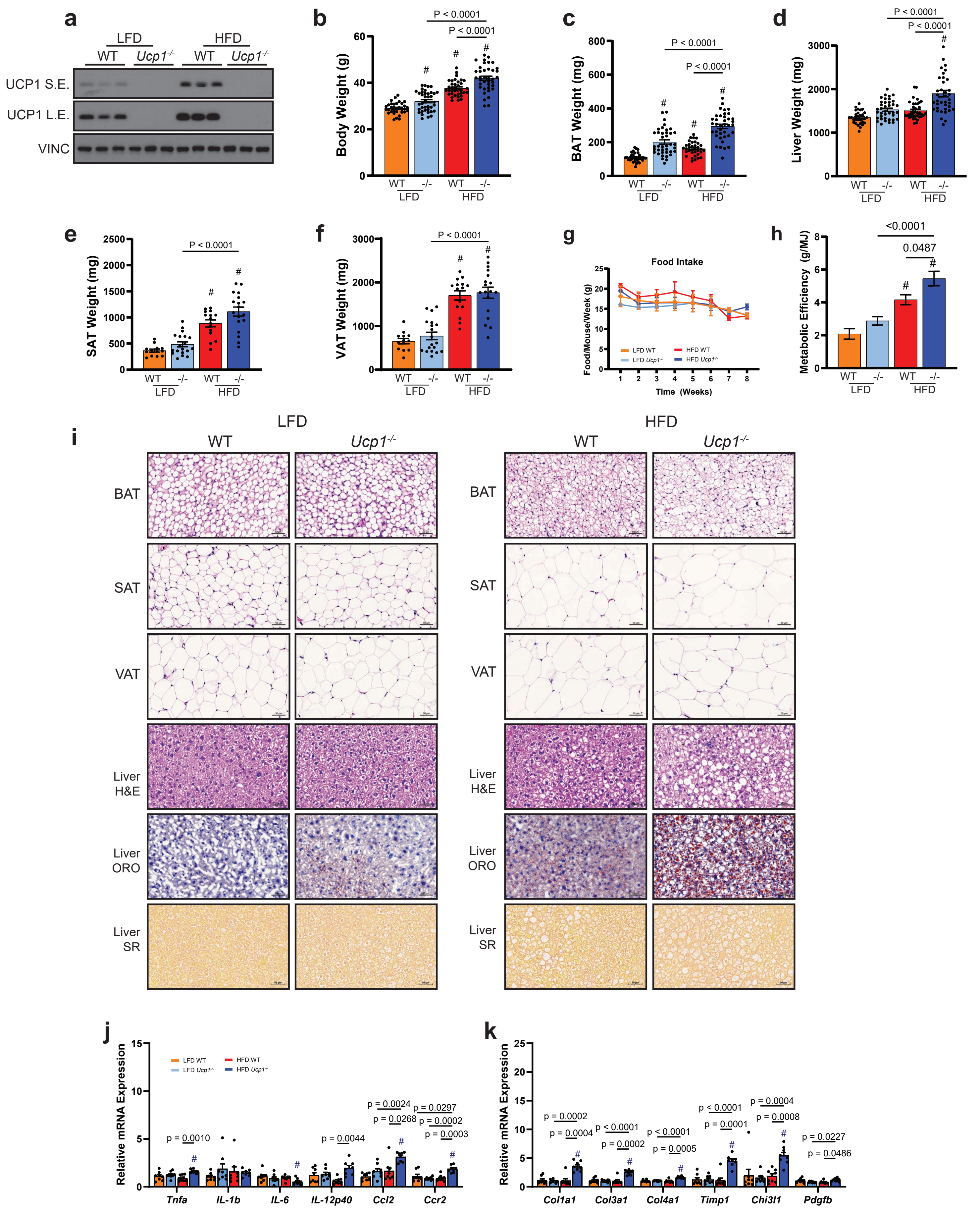


Extended Data Figure 3
Supports Figure 4

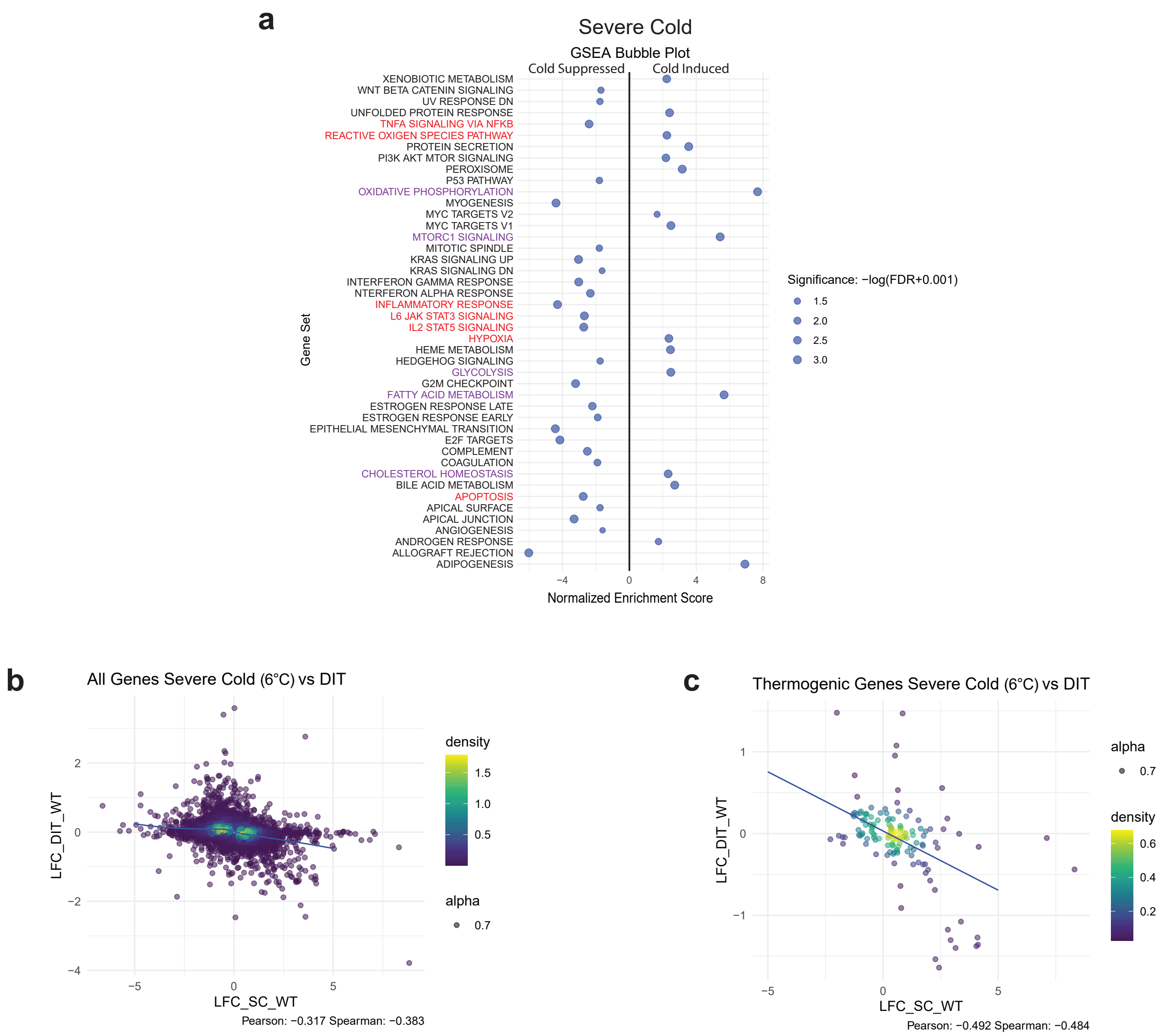


Extended Data Figure 4
Supports Figure 4

Extended Data Figure 5
Supports Figure 4

**a**

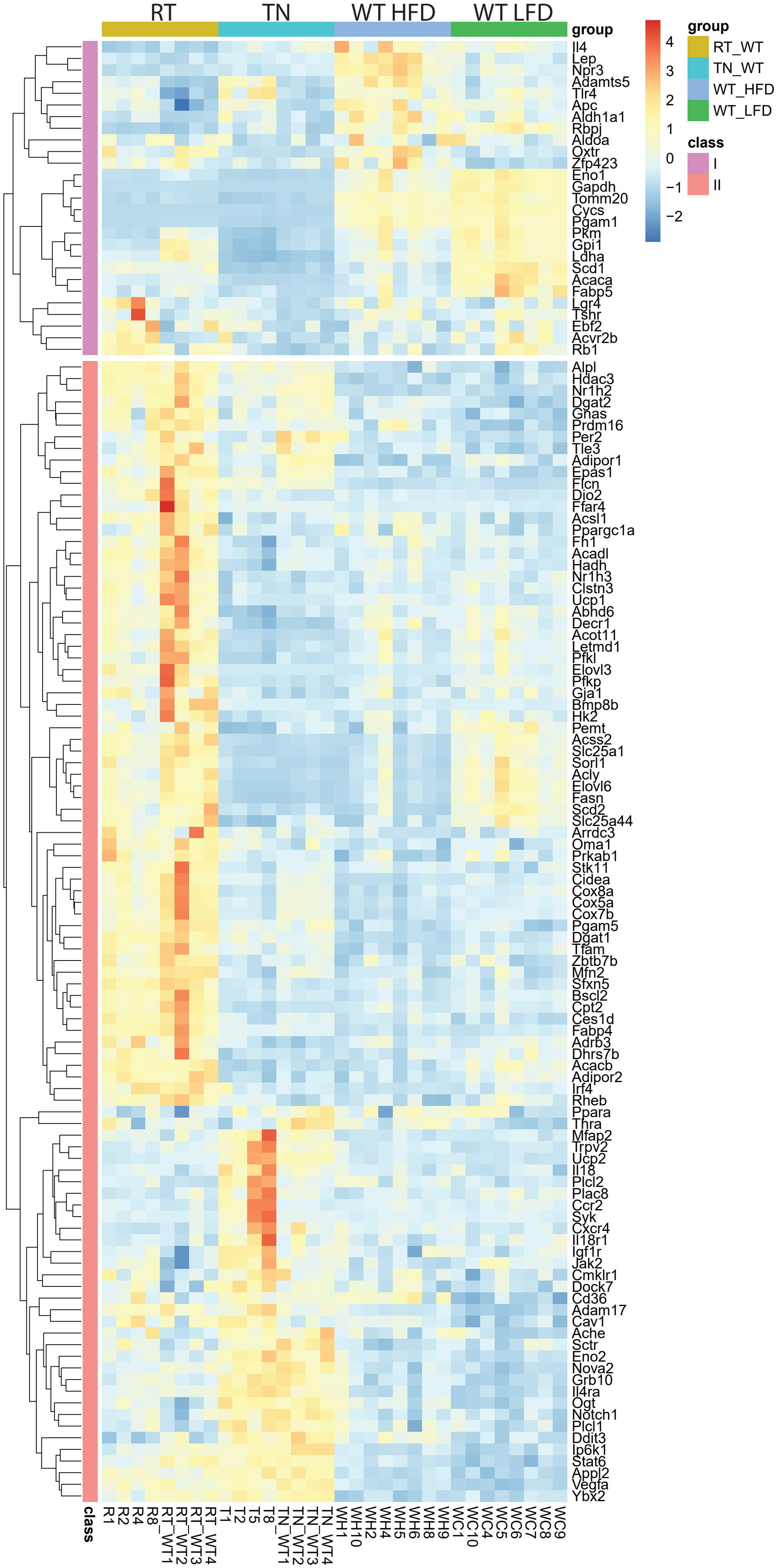

Extended Data Figure 6
Supports Figure 6

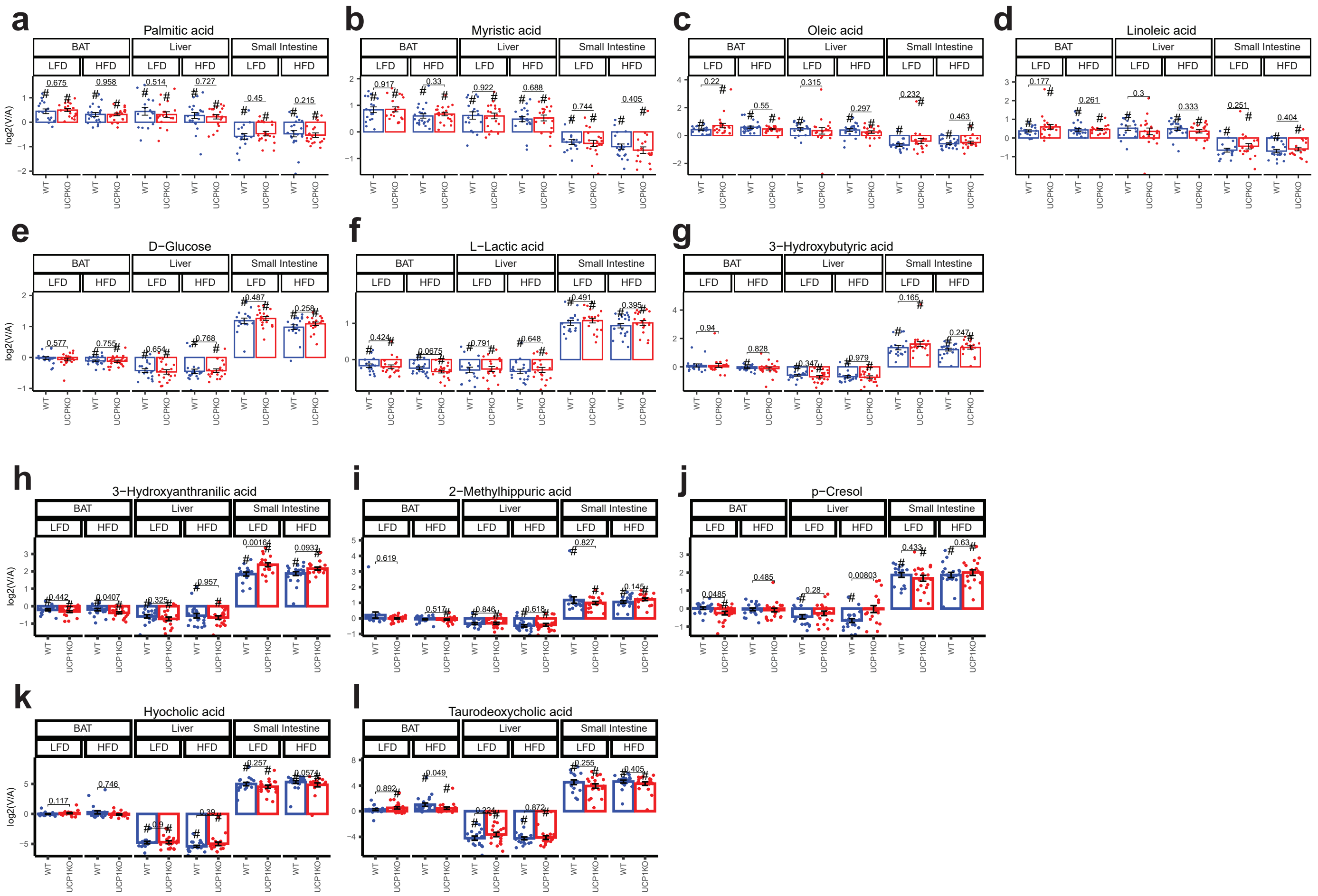

Extended Data Figure 7
Supports Figure 6

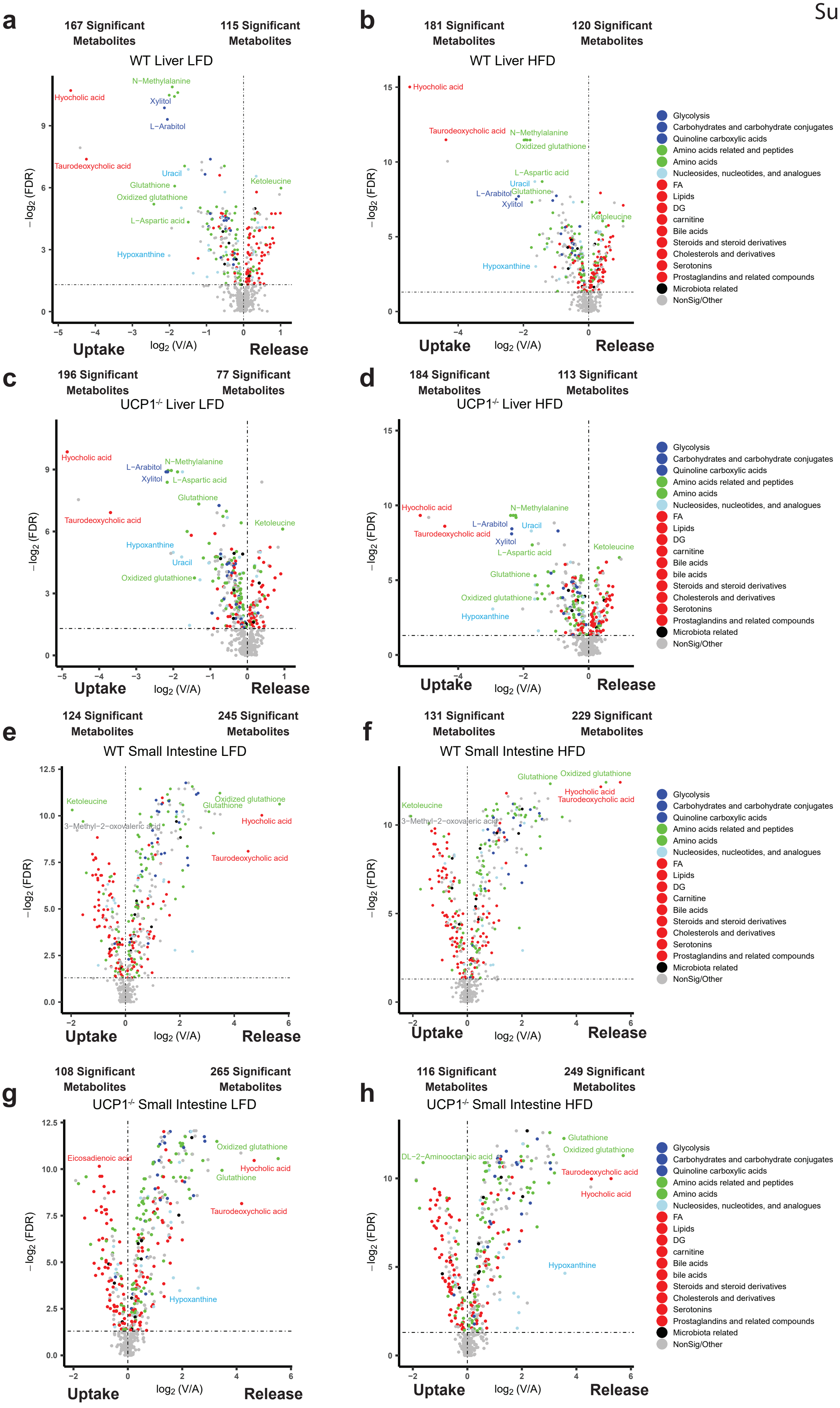

Extended Data Figure 8
Supports Figure 6

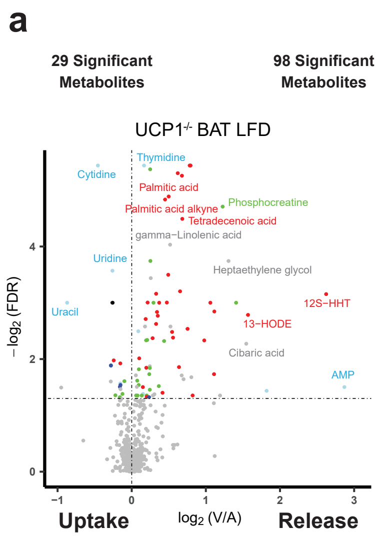


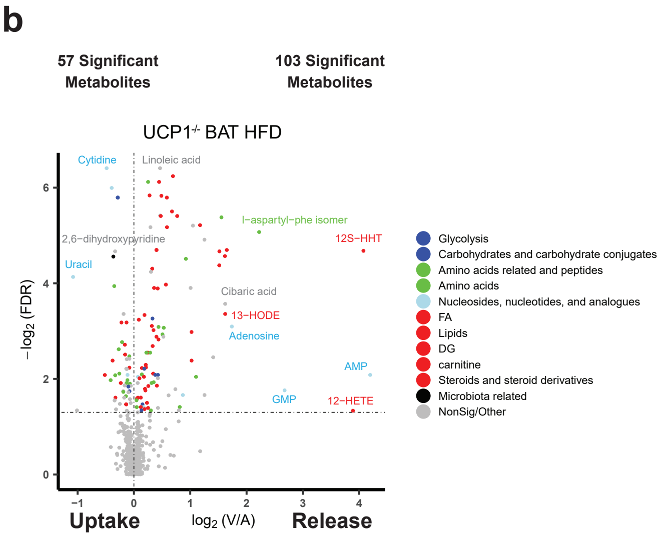


c

| | Median Blood Flow (ul/s per g BAT) |
|---|---|
| WT LFD | 23.10 |
| WT HFD | 19.45 |
| UCP1$^{-/-}$ LFD | 11.97 |
| UCP1$^{-/-}$ HFD | 8.33 |
